# A case study of *Whirly1* (*WHY1*) evolution in the angiosperms: altered sequence, splicing, and expression in a clade of early-transitional mycoheterotrophic orchids

**DOI:** 10.1101/2023.06.21.545690

**Authors:** Rachel M. Muti, Craig F. Barrett, Brandon T. Sinn

## Abstract

The plastid-targeted transcription factor *Whirly1* (*WHY1*) has been implicated in chloroplast biogenesis, plastid genome stability, and fungal defense response, which together represent characteristics of interest for the study of autotrophic losses across the angiosperms. While gene loss in the plastid and nuclear genomes has been well studied in mycoheterotrophic plants, the evolution of the molecular mechanisms impacting genome stability are completely unknown. Here we characterize the evolution of *WHY1* in four early-transitional mycoheterotrophic orchid species in the genus *Corallorhiza* by synthesizing the results of phylogenetic, transcriptomic, and comparative genomic analyses with *WHY1* genomic sequences sampled from 21 orders of angiosperms. We found an increased number of non-canonical *WHY1* isoforms assembled from all but the greenest *Corallorhiza* species, including intron retention in some isoforms. Within *Corallorhiza*, phylotranscriptomic analyses revealed the presence of tissue-specific differential expression of *WHY1* in only the most photosynthetically capable species and a coincident increase in the number of non-canonical *WHY1* isoforms assembled from fully mycoheterotrophic species. Gene- and codon-level tests of *WHY1* selective regimes did not infer significant signal of either relaxed selection or episodic diversifying selection in *Corallorhiza*, but did so for relaxed selection in the late-stage full mycoheterotrophic orchids *Epipogium aphyllum* and *Gastrodia elata*. Taken together, this study provides the most comprehensive view of *WHY1* evolution in angiosperms to date. Our analyses reveal that splicing alteration and decreased expression of *WHY1* are coincident with deceased plastome stability in a group of early-transitional mycoheterotrophic orchids and that these changes may precede the selective shifts observed in late-stage mycoheterotrophic species.

## 1 Introduction

The ability to photosynthesize has been lost dozens of times across the angiosperm Tree of Life, and at least 30 independent losses have occurred in the Orchidaceae (Merckx and Freudenstein, 2010; Barrett et al., 2014; Barrett et al., 2019). Mycoheterotrophy, the derivation of carbon nutrition from fungi (Leake, 1994), is common to all orchids during early development and is a nutritional requirement due to the lack of endosperm in their seeds (Rasmussen, 1995; 2002). In lieu of stored nutrition, orchid seeds have evolved a complex symbiotic relationship with fungi, where orchid seeds germinate only in the presence of an appropriate fungal partner and the nutrition required for embryo development is derived exclusively via the degradation of fungal hyphae which penetrate the orchid cells (Merckx, 2013; Zeng et al., 2016; Yuan et al., 2018). The duration of reliance upon their fungal partner for nutrition has been extended in some orchid species, which have evolved to parasitize fungi for the entirety of their lives. Independent shifts to a mycoheterotrophic condition throughout the angiosperms have independently led to plastid genome (plastome) degradation (Barrett et al., 2014; Wickett et al., 2016; Graham et al., 2017; Timilsena, et al., 2023), elevated rates of nucleotide substitution (Lemaire et al., 2010; Wicke et al., 2016), and oftentimes the loss of morphological structures such as leaves and roots (Leake, 1994).

*Corallorhiza* is a North American, temperate genus comprising 12 species of morphologically-reduced, mycoheterotrophic orchids for which varying states of plastid genome (plastome) degradation and inferred photosynthetic ability have been characterized (Zimmer et al., 2008; Cameron et al., 2009; Barrett and Freudenstein, 2008; Barrett and Davis, 2012; Barrett et al., 2014; Barrett et al., 2018). The presence of relatively intact plastomes containing the expected repertoire of housekeeping genes in *Corallorhiza* species evidences the clade as a group of early-transitional mycoheterotrophs, in contrast with late-stage mycoheterotrophic species which have highly degraded plastomes and lack many or all plastid housekeeping genes, such as *Epipogium* and *Gastrodia* species (*sensu* Barrett et al., 2012; see also Barrett et al., 2014). *Corallorhiza* species parasitize Basidiomycete fungi that are engaged in mycorrhizal relationships with nearby autotrophic plants, predominantly in the families Russulaceae and Thelephoraceae (Taylor and Bruns, 1997; Taylor et al., 2004; Barrett et al., 2010; Freudenstein and Barrett, 2014). Besides their relationship with fungi, a conspicuous characteristic of *Corallorhiza* species is the complete loss of both leaf laminae and roots. A recurrent theme of morphological reduction has been documented across parasitic and mycoheterotrophic plant lineages (see Leake, 1994), and work has recently focused on genomic content and gene expression in mycoheterotrophic species in order to improve our understanding of the genomic precursors and consequences of this trophic transition (Barrett et al., 2014; Wickett et al., 2016; Graham et al., 2017; Zhang et al., 2017; Yuan et al., 2018; Cai, 2023; Timilsena et al., 2023).

The integrity of the plastome of parasitic and mycoheterotrophic plants is of particular interest as reduction in gene content, increased number of pseudogenes, structural variation, and a reduction in overall genome length have been found to correlate with the degree of external carbon reliance among parasitic angiosperms. Generally, disruptions to the genome such as double-strand DNA breaks are harmful to the organism, and many mechanisms that help to protect against such occurrences have evolved throughout the Tree of Life (Waterworth et al., 2011). One such genome stabilizing mechanism that is increasingly recognized for its involvement in processes such as plastome double-strand break repair is the activity of the Whirly family of transcription factors (*WHY*; Desveaux et al., 2005).

Transcription factors are regulatory gene products that function by binding to DNA (Latchman, 1993). The Whirly family comprises *WHY1*, *WHY2*, and *WHY3*, which are three plant-specific, nuclear-encoded genes with DNA binding domains that are named for their whirligig-like structural conformation (Desveaux et al., 2002; Cappadocia et al., 2010). Crystal structures of the Whirly transcription factors have been determined as tetramers that have a single-stranded DNA binding domain that spans two subunits (Cappadocia et al., 2013). Of particular interest is *WHY1*, the product of which has been implicated to play roles in myriad processes including mediation of abiotic stressors (Zhuang et al., 2019, 2020; Ruan et al., 2022), induction of double strand DNA break repair (Cappadocia et al., 2010), and plastid biogenesis (Prikryl et al., 2008). Transcription factors are crucial to myriad regulatory and developmental processes, which is reflected in the massive expansions of many TF gene families in plants (Lehti-Shiu et al., 2017). Our present understanding of TF evolution has largely been informed through the investigations focused on understanding their roles in morphological or ecological diversification (see de Mendoza et al., 2013; Lai et al., 2020), rather than how they change in systems which have undergone coincident extreme loss of morphological and genomic features.

*WHY1* has been shown to dually localize to both plastids and the nucleus (Krause et al., 2005; Grabowski et al., 2008; Prikryl et al., 2008; Isemer et al., 2012; Ren et al., 2017). In chloroplasts, *WHY1* localizes to the boundary between the thylakoid and nucleoid membrane in chloroplasts and has been implicated in retrograde signaling regulating H_2_O_2_ homeostasis and as a coordinator of photosynthetic gene expression (Lepage et al., 2013; Foyer, et al., 2014; Lin et al., 2019). Species that have undergone the transition to heterotrophy experience elevated levels of oxidative stress compared to autotrophic relatives (Suetsugu et al., 2017; Lallemand et al., 2019). Additionally, *WHY1* proteins stabilize plastid genomes by non-specific binding to the genome which protects against microhomology-mediated DNA rearrangements, including deletions and duplications of sequences (Maréchal et al., 2009; Lepage et al., 2013; Zampini et al., 2015).

*WHY1* is involved in complex roles in both plant defense responses and genomic stabilization. It has long been known that mutations which reduce the binding affinity of *WHY1* correlate with increased infection by some pathogens (Desveaux et al., 2004). *WHY1* has been shown to bind to specific DNA promoter regions that can induce transcription (Zhuang et al., 2020), aid in defense response signaling and accumulation of disease resistance (Desveaux et al., 2004; Desveaux et al., 2005), and also maintain telomere length in the nuclear genome (Desveaux et al., 2005). Recently, *WHY1* has even been shown capable of negatively regulating the RNA interference response to two geminiviruses (Sun et al., 2023). Taken together, the literature suggests that modulation of *WHY1* expression can result in tradeoffs between plant defense and genomic stability.

The roles *WHY1* plays in stabilizing both the nuclear and plastid genomes (Yoo et al., 2007; Zampini et al., 2015), defense response (Desveaux et al., 2005), and chloroplast development (Qiu et al., 2022) are central to our choice to study the evolution of this transcription factor. In particular, experimental work demonstrating plastome destabilization (Lepage et al., 2013; Zampini et al., 2015), albinism, and variegation in *WHY1* mutants (Prikryl et al., 2008; Ren et al., 2017; Qiu et al., 2022) makes the gene a compelling target given the reduction in plastome content and length observed across parasitic angiosperms (Barrett et al., 2014; Wicke et al., 2016). To date, work characterizing the sequence and expression diversity of *WHY1* has been restricted to model or agricultural systems (Cappadocia et al., 2013; Ruan et al., 2022; Taylor et al., 2022) and no phylo-comparative investigations of sequence evolution and selective regime have been conducted. The implication of *WHY1* in processes associated with the mycoheterotrophic condition frames a phylogenetically-informed investigation of *WHY1* along a trophic gradient as an important step in improving our understanding of the evolution of plastid-targeted TFs in heterotrophic plant lineages.

We focus on four species, *C*. *trifida*, *C. striata*, *C. wisteriana*, and *C. maculata*, which together comprise an early-transitional trophic gradient to full mycoheterotrophy with sister relationships among partial and full mycoheterotrophs (Figure 1). The well-characterized phylogenetic relationships between these four *Corallorhiza* species (Barrett et al., 2018) provide a powerful framework upon which to investigate *WHY1* evolution during the early stages of transition to full mycoheterotrophy while accounting for phylogenetic non-independence (*sensu* Felsenstein, 1987). We consider *C*. *trifida* and *C. wisteriana* to be partial mycoheterotrophs, as their tissues contain measurable chlorophyll content and their plastomes are the most genetically-intact plastid genomes in the genus, although photosynthesis has only been directly observed in *C*. *trifida* (Zimmer et al., 2008; Cameron et al., 2009; Barrett et al., 2014; Barrett et al., 2018). Conversely, we consider *C. maculata* and *C. striata* to be fully mycoheterotrophic, evidenced by their highly reduced chlorophyll content and degradation of many photosynthesis-related genes (Barrett et al., 2014; Barrett et al., 2018).

**Figure 1.**
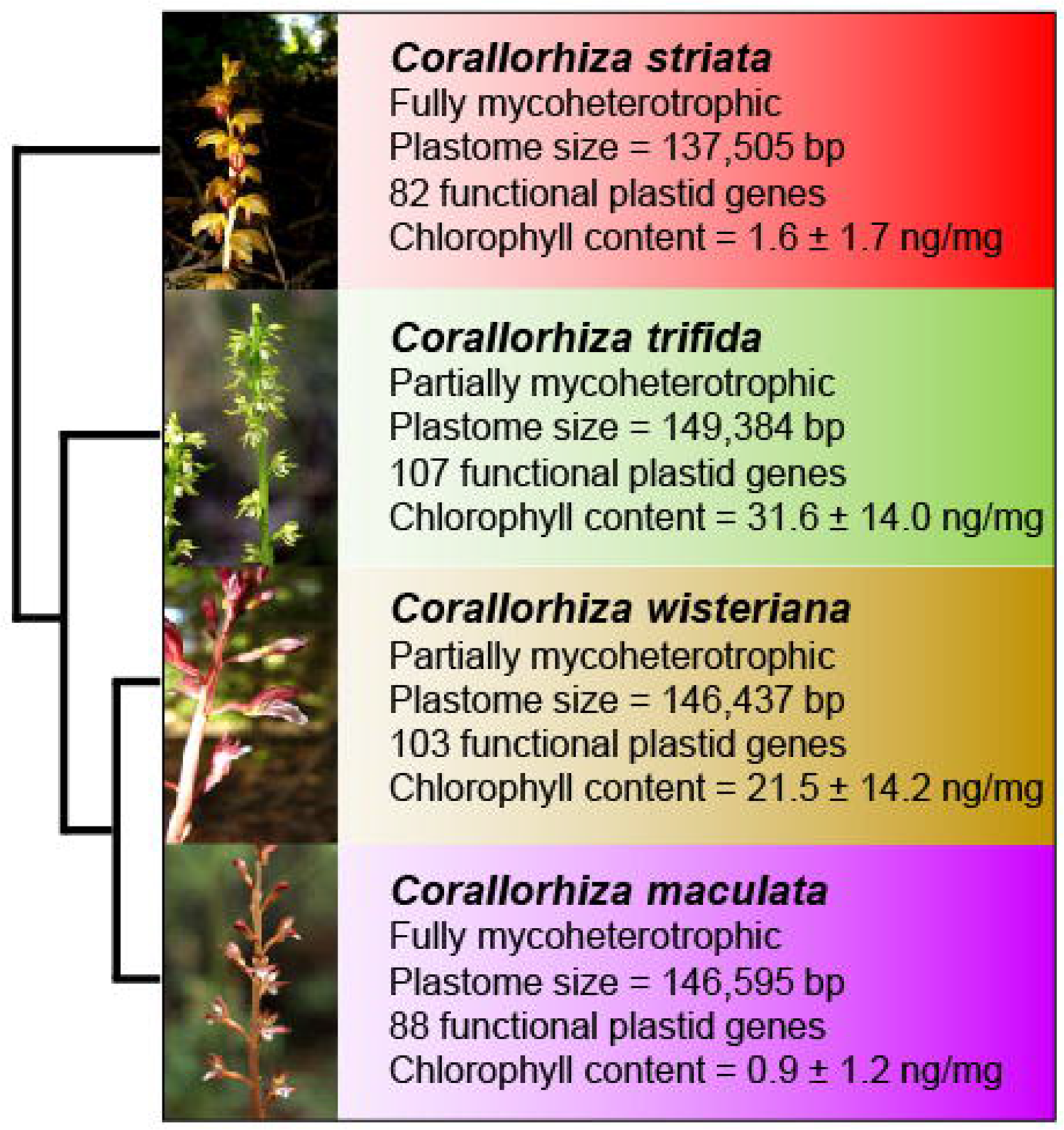
Overview of *Corallorhiza* species included in this study, showing plastid and nuclear phylogenetic relationships, inflorescence, trophic status (fully vs partially mycoheterotrophic), plastome size (bp), number of putatively functional plastid genes, and chlorophyll content (mean and standard deviation in nanograms of total chlorophylls per milligram of plant material). Phylogenetic, plastome and chlorophyll content data are from Barrett et al. (2014).

Here we characterize the evolution of *WHY1* across a mycoheterotrophic gradient, framed by phylogenetic context provided by the broadest taxonomic sampling of the gene to date, including late-stage mycoheterotrophic orchid species from the genera *Epipogium* and *Gastrodia*. Our integrative work leverages a combination of novel and publicly available data generated from DNA sequencing, RNAseq, and Oxford Nanopore sequencing from 110 angiosperm species representing 21 orders. Taken together, the results of our analyses of *WHY1* sequence, expression, splicing, and selective regime across both a trophic gradient and the angiosperms more broadly suggest that the gene may play a critical role in maintaining plastome stability after the transition to mycoheterotrophy.

## 2 Materials and Methods

### Publicly-available Sequences

Annotated *WHY1* sequences were obtained from the Nucleotide and RefSeq NCBI databases and Orchidstra, an orchid-specific database (Chao et al., 2017). Only *WHY1* sequences containing canonical *WHY1* ORFs (open reading frames) were retained. Additionally, sequences were excluded if they did not contain the ssDNA binding region (KGKAAL; *A. thaliana* Q9M9S3) as reported by Cappadocia et al. (2013; PDB 4KOO). Identical sequences were excluded for taxa with multiple database accessions. Stop codons were trimmed from sequences, with the exception of premature stop codons in sequences from known mycoheterotrophs, which were changed to gap characters (-) for compatibility with downstream methods (e.g. HyPhy, see below). In total, *WHY1* sequences from 110 species were included in the angiosperm-wide alignment (see Alignment section below; Table S1).

### RNAseq and *De Novo* Assembly of Transcripts

*Corallorhiza* tissues used for RNAseq are those referred to in Sinn and Barrett (2020). In brief, tissues were categorized as either aboveground (combined stem, flower, and ovary tissues) or belowground (rhizome tissue, including fungal tissue), and three biological replicates of each tissue type were sequenced for all four species, with the exception of aboveground *C*. *maculata* (two samples) and belowground *C*. *striata* (two samples) for which library preparation of limited material was not successful. *De novo* assembly of transcripts from each *Corallorhiza* species was conducted using Trinity (version v2.13.2, Grabherr et al., 2011). Reads were trimmed using Trimmomatic (version 0.36, Bolger et al., 2014) with default settings to remove sequencing adapters and low-quality bases from the read ends. Trinity was provided with a samples file with biological replicate relationships, and SS library type was set to RF. Trimmed reads were mapped to each assembled transcript using the splice-aware read mapper BBMAP (version 38.96; Bushnell, 2022) with default settings, with mapped reads output to SAM format and converted to sorted BAM-formatted files using SAMtools (version 1.15; Li et al., 2009).

### Identification of *WHY1* Transcripts

We used the HMMER suite (version 3.3.2; Eddy, 2011) to create a *WHY1* hidden markov model (HMM) profile. The HMM profile was built with hmmbuild, which used the complete angiosperm-wide *WHY1* nucleotide alignment. All assembled *Corallorhiza* transcripts were then searched against the HMM model using nhmmer (Wheeler and Eddy, 2013). Default parameters were used for both programs. All assembled isoforms of a transcript identified with the highest E-value were considered as potential splicing variants of *WHY1*.

### PCR Amplification

Primers for amplification of the entire *WHY1* sequence (Table S2) were designed using the Geneious Prime (version 2020.2.4, https://www.geneious.com) plugin for Primer3 (version 2.3.7; Untergasser et al., 2012). Oligos were synthesized by Integrated DNA Technologies (Coralville, Iowa, USA). PCR amplification of *WHY1* for each *Corallorhiza* species was performed using the Taq RED PCR Kit from Bio-Rad Industries (Hercules, California, USA). PCR was conducted in a BioRad T100 Thermocycler using the following program: initial template denaturation at 95°C for three minutes, followed by 30 cycles of denaturation at 95°C for 30 seconds, primer annealing at 52°C for 30 seconds, and template extension at 72°C for 30 seconds. The program ended with a final extension at 72°C for five minutes and was held at 4°C until retrieval. PCR clean-up was performed with Agencourt AMPure XP PCR Purification beads (Beckman Coulter Life Sciences; Indianapolis, Indiana, USA). Purified DNA samples were quantified using a NanoDrop One^C^ spectrophotometer (ThermoFisher Scientific; Waltham, Massachusetts, USA).

### Nanopore Sequencing

Purified DNA samples were diluted to equimolar concentrations and libraries for long-read sequencing were prepared according to the Oxford Nanopore Technologies (ONT; Oxford, United Kingdom) End-Prep protocol (SQK-LSK109). The library of each *Corallorhiza* species received a unique barcode for Nanopore sequencing using the Native Barcoding Expansion 1-12, PCR-free kit (EXP-NBD104). The MinION SpotON flow cell (R9.4.1 FLO-MIN 106; ONT) was used for sequencing. Basecalling was performed using the high-accuracy base calling algorithm as implemented in the GPU version of Guppy (version 6.2.1+6588110a6; ONT) on an NVIDIA GeForce RTX 2060 graphics card. Nanopore reads were mapped to the *Dendrobium catenatum WHY1* genomic sequence in Geneious Prime using Minimap2 (version 2.17, Li, 2018) with a K-Mer length set to 15.

### Alignment

We aligned all *Corallorhiza WHY1* isoforms using MAFFT (version 7.3.10; Katoh and Standley, 2013), and the E-INS-i algorithm (Katoh et al., 2002) and the 1PAM scoring matrix, as a Geneious Prime plugin. Visualization and structural annotation against the canonical *WHY1* sequence of *Arabidopsis thaliana* (NCBI Q9M9S3) was also conducted using Geneious Prime. Consensus was determined as majority consensus with a 0% threshold, meaning no minimum frequency was required for a consensus character if the character was shared by most sequences. All *Corallorhiza WHY1* isoforms were translated in Geneious Prime to amino acid sequence and manually trimmed to the correct ORF. ORF-trimmed *WHY1* translations were aligned with the ORF-trimmed *WHY1* sequence of *Arabidopsis thaliana* (NCBI NM101308) to verify the presence of the expected canonical reading frame.

A translation-aware alignment of the canonical form of *WHY1* representing lineages across the angiosperms was generated using a two-step process. Nucleotides were first aligned and translated using the frameshift-aware aligner MASCE2 (version 2.0.6; Ranwez et al., 2011) with default parameters, which inserts gap characters necessary to maintain codon-based statements of homology across the alignment. This approach was necessary due to the presence of frameshift mutations in sequences from *Gastrodia elata*, *Epipogium aphyllum*, and *C*. *striata*. The MACSE2-processed nucleotide and amino acid alignments were then refined using MAFFT and the E-INS-i algorithm with a BLOSUM 80 substitution matrix.

An alignment including the complete genomic sequences of *WHY1* from *Phalaenopsis equestris* (ASM126359v1) and *Dendrobium catenatum* (ASM160598v2) was also generated. *Phalaenopsis equestris* (Cai et al., 2015) and *D*. *catenatum* (Zhang et al., 2016) are the closest relatives of *Corallorhiza* with sequenced genomes. This DNA alignment was generated to identify the introns of *WHY1* and to evaluate the exonic content of assembled transcripts. All *Corallorhiza* isoforms and *Corallorhiza* Nanopore consensus sequences were aligned with the sequences of *P*. *equestris* and *D*. *catenatum* using MAFFT (version 7.3.10; Katoh and Standley, 2013), as a Geneious Prime plugin, and the E-INS-i algorithm (Katoh et al., 2002).

### Visual Screening for Amino Acid Substitutions

Nonsynonymous substitutions within the angiosperm *WHY1* alignment were surveyed visually in our amino acid alignments via Geneious Prime. Substitutions of interest included those present exclusively in *Corallorhiza* species, mycoheterotrophs, and the Orchidaceae. The codons of *A*. *thaliana* (Q9M9S3) and *P*. *equestris* that corresponded to the sites of nonsynonymous substitutions in *WHY1* of interest were cross-referenced with the annotated *A*. *thaliana WHY1* sequence for structure and the *P*. *equestris WHY1* genomic sequence as included in the DNA alignment for exon location.

### Phylogenetic Inference

IQ-TREE (version 1.6.12; Nguyen et al., 2015) was used to infer phylogenetic relationships amongst the recovered *WHY1* sequences using automated model choice (Kalyaanamoorthy et al., 2017), optimal partitioning assessment, and nearest neighbor interchange search was enabled. Node support was estimated using 1,000 ultrafast bootstrap approximation replicates (Hoang et al., 2018). Two partitions were defined, which comprise the signal peptide and the highly variable 5’ portion of the chain (positions 1 – 516) and the highly conserved chain region (517 – 1,122), identified using functional annotations on *WHY1* sequence per *A. thaliana* (Q9M9S3).

### Differential Expression

*In silico* analysis of differential expression was conducted using scripts provided as components of the Trinity RNAseq pipeline (Haas et al., 2013). Transcript abundance was estimated using the alignment-free estimation method as implemented in Salmon (version 1.2.0, Patro et al., 2017). The strand-specific library type was set to RF, for reverse-forward. Salmon output files included transcript abundance estimates at both the transcript and gene level. Matrices were built using the abundance_estimates_to_matrix.pl script for both transcript counts and gene expression. Differential expression analysis was run using the R (R Core Team, 2019) package edgeR (Bioconductor version 3.10, Robinson et al., 2010) via the run_DE_analysis.pl script, which identified differential expression using biological replicates of tissues across the replicate conditions aboveground and belowground. TPM (transcripts per million) values were normalized using the TMM (trimmed mean of M-values; Robinson and Oshlack, 2010) approach in order to normalize expression values while maintaining comparability of expression between samples.

### Selection Analyses

The *WHY1* nucleotide alignment was tested for statistically significant changes in selection regime using four methods implemented in the command-line, multithreaded version of the Hypothesis Testing using Phylogenies suite (HyPhy; version 2.5.39; Pond and Muse, 2005). We tested for significant change of selective regime using five test branch sets against null reference branch sets comprising all other species in the phylogram: 1) *Corallorhiza* species; 2) *C*. *maculata* + *C*. *striata*; 3) *Corallorhiza* + *Epipogium aphyllum + Gastrodia elata*; 4) *C*. *maculata* + *C*. *striata* + *E*. *aphyllum + G*. *elata*; 5) *E*. *aphyllum* + *G*. *elata*. The same *Amborella trichopoda*-rooted maximum likelihood *WHY1* topology inferred with IQ-tree was used for all analyses.

RELAX (Wertheim et al., 2015) was used to test for evidence of relaxed selection. RELAX breaks each codon into its three component sites, each with an assigned omega class. Values for omega are calculated as Dn/Ds ratios, using the calculation for the reference branches as the null hypothesis. The value k is the selection intensity parameter and is an exponent value on omega. The alternative model fits a value for k that changes the rate to fit with the test branches. Evidence for intensified selection strength along test branches is indicated by a significant result where the value of k is greater than 1 (k > 1, P < 0.05). Evidence for relaxed selection along test branches is indicated by a significant value of k less than 1 (k < 1, P < 0.05). Strength of selection was assessed simultaneously for all species using Fast Unconstrained Bayesian Approximation (FUBAR, Murrell et al., 2013). FUBAR can detect weak, yet pervasive, purifying or diversifying selection at the codon level (posterior probability > 0.90) without the use of test and reference branch sets.

The Branch-Site Unrestricted Statistical Test for Episodic Diversification (BUSTED) was used to test for evidence of gene-wide positive selection (Murrell et al., 2015). BUSTED uses three omega classes defined as ω1≤ω2≤1≤ω3. The ω1 class is the proportion of sites with a very low Dn/Ds ratio. The ω2 class is the proportion of sites just below 1, and ω3 are sites above 1. A value of 1 suggests selective neutrality, and therefore defines the null, or Constrained, model. BUSTED then calculates the log-likelihood of the data for each of the null and alternative models. These ratios are calculated for each site and are called evidence ratios (ERs). They are used as a threshold (χ^2^ distribution, P < 0.01) but are not a valid test for site-specific likelihood. The null model is rejected if at least one site on a test branch experienced positive selection. Evidence for positive, or diversifying, selection in a gene of a test branch is indicated by a rejection of the null model.

The Adaptive branch-site random effects likelihood (aBSREL) test was used to test for signal of episodic positive selection. aBSREL infers ω values at both the level of sites and branches and can account for rate heterogeneity inherent to complex evolutionary scenarios by partitioning these values into multiple rate classes per branch. aBSREL was conducted in exploratory mode, where all branches were tested and p-values were Holm-Bonferroni corrected, and in the more sensitive *a priori* mode to test for episodic positive selection in each test branch set.

## 3 Results

### *De novo* Assembly and Identification of *WHY1*

The hmmer suite revealed a single, Trinity-identified gene model and its isoforms as representing *WHY1* from *de novo* assembled transcripts for each *Corallorhiza* species. Trinity assembled a single isoform representing the expected canonical CDS (coding DNA sequence) of *WHY1* for all but *C. striata*, the latest-stage fully-mycoheterotrophic species of *Corallorhiza* sampled. However, mapping reads to each isoform with BBMAP revealed that a five nucleotide indel, the absence of which results in a premature stop codon, was differentially present or absent in the reads of each *Corallorhiza* species (NCBI BioProject PRJNA984634). Trinity differentially incorporated that indel, hereafter referred to as the GTGAA indel, into the isoform pool of each *Corallorhiza* species. The absence of the GTGAA indel in Trinity-assembled isoforms of *C*. *striata* precluded the recovery of the canonical ORF, but read mapping supports the presence of the five nucleotides necessary to recover the expected ORF in *C*. *striata*, at a rate of 67.2% of reads in isoform 2 and 76.1% of reads in isoform 4. Those data support that the canonical variant of *WHY1* is also expressed. We analyzed the *C*. *striata* isoforms as assembled, rather than manually modifying the *C*. *striata* transcripts to conform to a hypothesized canonical sequence. The inclusion of those five nucleotides would not alter any results presented here, aside from whether a canonical *WHY1* isoform is transcribed in *C*. *striata*.

The canonical *WHY1* ORF in *C*. *trifida*, *C. wisteriana*, and *C. maculata* is 795 nucleotides in length and comprises 265 amino acids, two fewer amino acids than that of *A*. *thaliana*. Non-canonical splicing variants were found in the isoform pools of each *Corallorhiza* species. A splicing variant containing a 79-nucleotide-long sequence of 96.6% mean pairwise identity was recovered from *C*. *wisteriana*, *maculata*, and *striata*, at nucleotide position 232. The second major variant is the GTGAA indel discussed earlier, which is variously present at nucleotide position 514 in all four species. A third variant identified from the isoform pool of *C*. *striata* represents a modification of the 3’ end of the ORF, with sequence of no obvious homology to that of *WHY1* immediately downstream of the previously discussed five-nucleotide indel that was variously present in our read pools throughout *Corallorhiza*.

### Alignment

MAFFT alignment of all raw Trinity-assembled *Corallorhiza WHY1* isoforms resulted in a matrix of 1,302 positions. Pairwise percent identity across isoforms was 81.4% and 88.2% and gaps comprised 11.7% and 8.1% of character states for all isoforms and with the exclusion of the early terminating isoform 1 of *C*. *striata*, respectively. Alignment of ORF-trimmed transcripts resulted in a matrix of 874 positions (Figure 2). Percent pairwise identity amongst isoforms within a species was highest for *C*. *trifida* (99.5%) and lowest for *C*. *striata* (70.6%). A 79-nucleotide-long indel of 96.6% pairwise identity was identified in at least one isoform assembled from all *Corallorhiza* species except *C*. *trifida*. Translation-aware alignment of ORF-trimmed canonical *Corallorhiza WHY1* sequences, with the differentially present GTGAA indel manually inserted into the otherwise canonical sequence of *C*. *striata* resulted in an alignment of 795 characters of 98.0% pairwise identity and no gaps. The highest pairwise percent identity was 99.37% between *C*. *wisteriana* and *maculata* and the lowest was 96.98% between *C*. *trifida* and *striata*.

**Figure 2.**
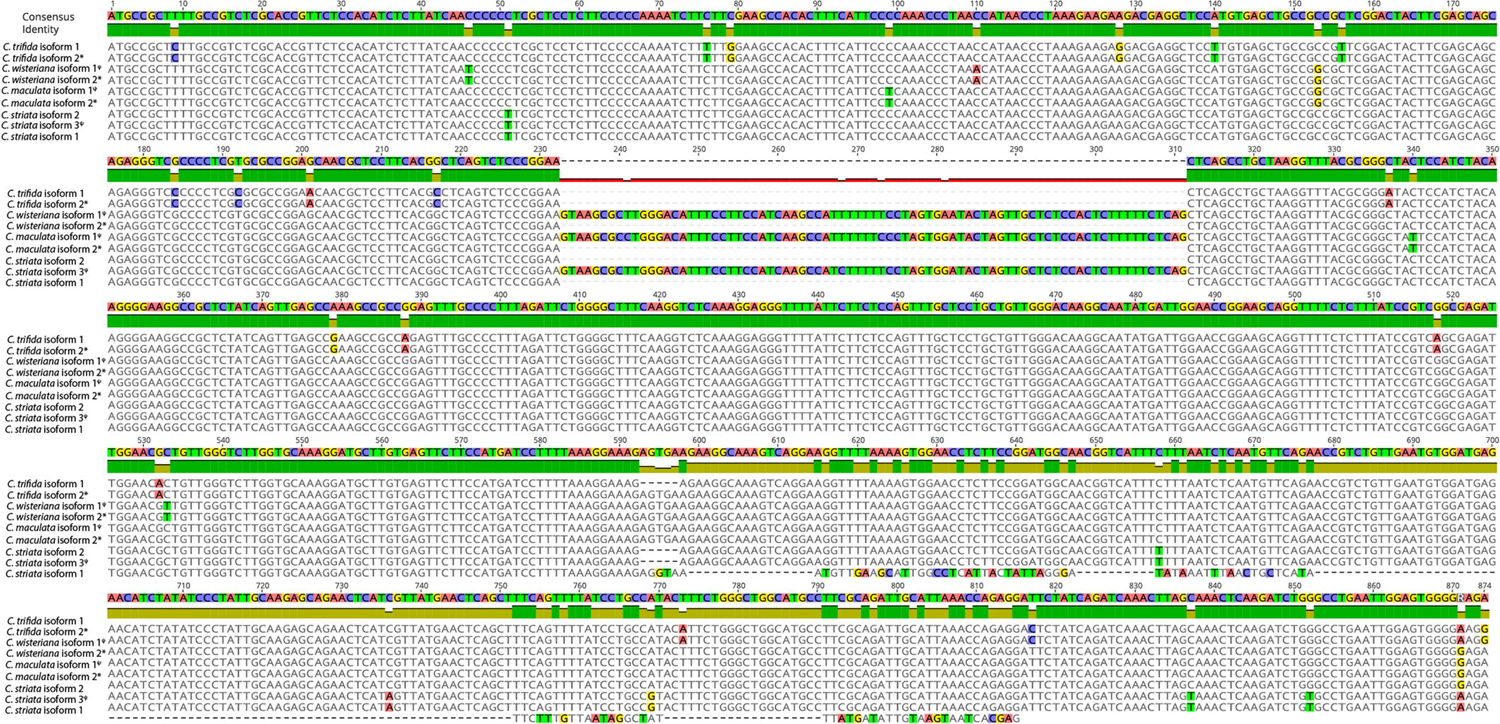
Wrapped view of open reading frame-trimmed *Corallorhiza WHY1* isoform alignment. Disagreements with the consensus sequence are highlighted. An * denotes a canonical isoform while ^ψ^ denotes an isoform with a retained intron.

The *WHY1* genomic DNA alignment of *Corallorhiza* isoforms, *Corallorhiza* Nanopore sequences, and *P. equestris* and *D. catenatum* genomic sequences had a total length of 12,000 nucleotides (Figure 3), in which the assembled canonical isoforms from each *Corallorhiza* species contain the expected exons of *WHY1*. One non-canonical isoform of *C*. *wisteriana*, *C*. *maculata*, and *C*. *striata* each contained intron 1, for which pairwise percent identity was 96.6% across those species. Percent pairwise identity of intron 1 between that of *D. catenatum* and *P*. *equestris* was 82.3%, and similarity between retained introns and that of *P*. *equestris* ranged from 78.5% to 82.3% for *C*. *maculata* and *striata*, respectively. The Nanopore-sequenced and Trinity-assembled *WHY1* intronic sequences of *C*. *striata* were identical, while those of *C. maculata* were 94.9% similar, due to the relative lack of two thiamine nucleotides in the Nanopore sequence.

**Figure 3.**
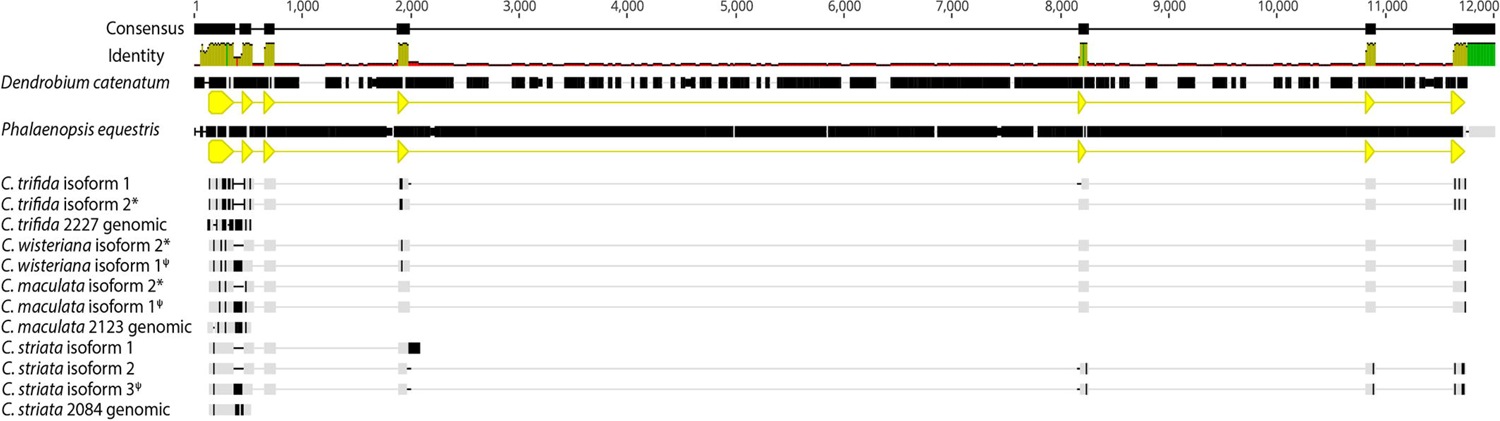
DNA alignment of canonical isoforms and Nanopore-sequenced genomic sequence from each *Corallorhiza species* with annotated *WHY1* sequence from the *Phalaenopsis equestris* and *Dendrobium catenatum* reference genomes. *WHY1* exons 1 and 2 are annotated in yellow and the locus ID tags for each reference genome are depicted in both the gene (green) and CDS (yellow) annotations. Disagreements with the consensus sequence are shown in black. Percent identity of aligned sites is depicted as a histogram. An * denotes a canonical isoform while ^ψ^ denotes an isoform with a retained intron.

The angiosperm-scale *WHY1* nucleotide alignment contained *WHY1* sequences from 110 species, including 22 orchid species. The total length of the alignment was 1,122 positions, of which 150 were of identical states across all species. Two partitions roughly corresponding to the transit peptide and chain regions as annotated in *A. thaliana* (Q9M9S3) were conspicuously visible in the consensus sequence of the alignment. The first was a highly variable, lineage-specific portion of sequence ranging from positions 1 – 516 in the alignment. The second was a highly conserved portion spanning positions 517 – 1,122. The mean pairwise percent identity of the transit region of Orchidaceae was 51.3%, while that of *Apostasia shenzhenica* compared to either *G*. *elata* or *E*. *aphyllum* was 21.5% and 31.1%, respectively. The mean pairwise percent identity of the transit region of *Corallorhiza* was 94.4%. The mean pairwise percent identity of the four *Corallorhiza* species in the alignment was 97.7%, and 81.0% across the Orchidaceae. The grasses had the lowest mean percent pairwise similarity of any clade, which was 49.36% when each sequence was compared to each non-grass species in the alignment.

The angiosperm-wide *WHY1* amino acid alignment comprised 376 positions, of which 48 were of identical states across all species. The two partitions in the nucleotide alignment corresponding to functional annotation in *A. thaliana* (Q9M9S3) were more conspicuous in the amino acid alignment. Positions 1 – 115 and 116 – 376 corresponded to the transit and chain regions of *A*. *thaliana* (Q9M9S3), respectively. The pairwise percent identity of the four *Corallorhiza* species in the alignment was 97.4% *WHY1*, and across the Orchidaceae it was 77.8%.

### Nanopore Sequencing

Nanopore sequencing of genomic *WHY1* sequence from three of the four *Corallorhiza* species confirmed that the unique sequence in the non-canonical transcripts of *C. wisteriana*, *C. striata*, and *C. maculata* represented retention of a *WHY1* intron 1. Sequencing of *WHY1* amplicons generated read pools ranging from 25,546 to 74,261 reads in *C*. *trifida* and *C*. *maculata*, respectively. Mapping of Nanopore reads against the *D*. *catenatum* genomic *WHY1* sequence resulted in a mean coverage depth for the exon 1-2 region ranging from 4,034.9 to 10,617.6 in *C*. *trifida* and *C*. *striata*, respectively (Table S2). Library preparation for Nanopore sequencing of *C*. *wisteriana WHY1* amplicons was deemed unsuccessful since the resulting sequence pool contained presumably off target reads that precluded confident assembly of a *WHY1* consensus sequence from that sample. Alignment of the Nanopore-sequenced *WHY1* amplicon consensus reads with the *Corallorhiza* transcript isoforms and the full *WHY1* genomic sequences of *P*. *equestris* and *D*. *catenatum* provided evidence that the non-canonical isoforms of *WHY1* in *Corallorhiza* were a result of alternative splicing (Figure 4). At least one non-canonical isoform from all but *C*. *trifida* contains intron 1 of *WHY1*.

**Figure 4.**
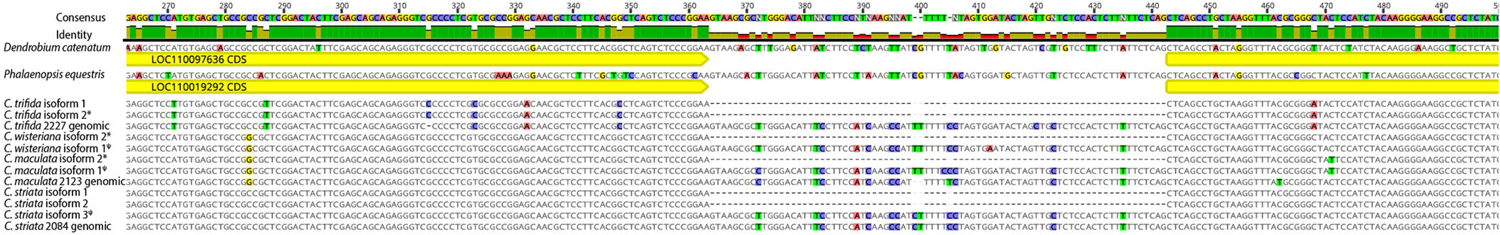
DNA alignment of *Corallorhiza* isoforms assembled from each *Corallorhiza species* and Nanopore sequences for *C*. *trifida*, *maculata*, and *striata* with annotated *WHY1* sequence from the *Phalaenopsis equestris* and *Dendrobium catenatum* reference genomes. *WHY1* exons 1 and 2 are annotated in yellow and the locus ID tags for each reference genome are depicted in both the gene (green) and CDS (yellow) annotations. Disagreements with the consensus sequence are highlighted. Percent identity of aligned sites is depicted as a histogram. An * denotes a canonical isoform while ^ψ^ denotes an isoform with a retained intron.

The sequences obtained from *Corallorhiza* species via Nanopore and RNAseq shared a high degree of similarity with each other and the *WHY1* sequence in the previously published *P. equestris* and *D. catenatum* genomes. Intron 1 of *WHY1* was found to have a pairwise percent similarity of 82.3% between *P. equestris* and *D. catenatum*. *Corallorhiza WHY1* intron 1 sequences obtained via Nanopore sequencing and RNAseq had a mean pairwise percent similarity of 96.9%, while that of *C*. *striata* obtained via both RNAseq and Nanopore sequencing had a pairwise percent similarity of 82.3% with *P. equestris*.

### Select Amino Acid Substitutions of Interest

Substitutions exclusive to mycoheterotrophic species were evident and in some cases were exclusive to fully-mycoheterotrophic species. The transit regions of both *E*. *aphyllum* and *G*. *elata* contained some substitutions that shared little homology with the remainder of Orchidaceae. A phenylalanine-glycine residue at positions 76-77 was exclusive to *C*. *trifida*, while a leucine-arginine residue was present for the remainder of Orchidaceae at those positions. While pairwise percent identity of positions 76-77 was only 13.3% throughout angiosperms, the leucine-arginine residue in the Orchidaceae had a pairwise percent identity of 90.9% when including the residue from *C*. *trifida* and was identical that taxon was excluded. Substitutions of single amino acids comprising the transit region not found in other orchids were identified in both *C*. *wisteriana* (positions 46, 95) and *C*. *trifida* (position 113), but identical amino acids were identified in some non-orchid autotrophs.

At least one substitution that could impact *WHY1* conformation was identified in *Corallorhiza*. The terminal codon of an alpha helix annotated in the structure of *A*. *thaliana* (9M9S3) *WHY1* (position 336) has been substituted to an isoleucine in *C*. *trifida* and to a phenylalanine in *C*. *wisteriana*, *maculata*, and *striata* (Figure 5). We infer that the ancestral state of position 336 is leucine, on the basis of that codon state for 89.1% of sampled taxa, including *Amborella trichopoda*. The only other substitutions at that position were to isoleucine, which was identified in eight autotrophic angiosperms outside the Orchidaceae, and in *C*. *trifida*. Additionally, a substitution from alanine to valine at position 359 was exclusive to *C*. *striata*, which represents the only occurrence of that state for this position for 109 other angiosperm species, and the only subgeneric polymorphism observed in that position.

**Figure 5.**
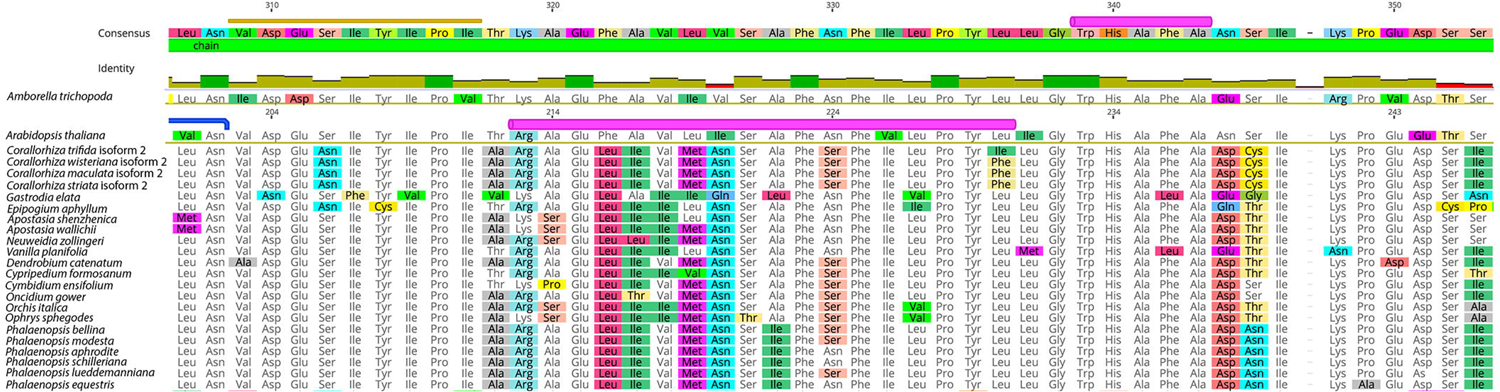
Detailed view of amino acid alignment of *WHY1* sequences of orchid species. Note the phenylalanine substitutions unique to *Corallorhiza wisteriana*, *maculata*, and *striata*, relative to the other species sampled. The amino acid substitutions of interest are located at the 3’ portion of an region inferred to conform into an alpha helix structure (pink annotation) in *Arabidopsis thaliana WHY1* (alignment row two).

A five-nucleotide indel resulting in a frameshift (amino acid alignment positions 269 & 270) was found in *C*. *striata*, but the pairwise percent identity of downstream sequence with that of *C*. *maculata* was high (97.8%) and reads containing the corresponding five nucleotides were identified in the RNAseq read pool. In fact, RNAseq reads containing the indel were found in the read pools of all *Corallorhiza* species, suggesting that isoform diversity was conservatively interpreted by our methods.

We found that the chain region of *E*. *aphyllum* contained a glycine to arginine substitution in the characteristic *WHY1* ssDNA binding motif (positions 188-193), a site that was otherwise conserved throughout the remainder of samples. Additionally, we found that the *E*. *aphyllum* sequence contained a residue comprising seven amino acids (positions 294 – 300), the last of which was a premature stop codon. MAFFT resolved those seven amino acids as an insertion with no homology to other angiosperm sequences. Six substitutions downstream of that premature stop codon are exclusive to *E*. *aphyllum*.

### Phylogenetic Inference

Both the Eudicots and Monocots were recovered as monophyletic (BS = 100%). Orchidaceae was recovered as a monophyletic group (BS = 94%), with *D. catenatum*, *G. elata* and *E. aphyllum* as an early-diverging paraphyletic grade within the Epidendroids (BS = 98%). *Corallorhiza* species were recovered as a monophyletic group (BS = 100%) with *C*. *trifida* sister to *C*. *striata* + (*C*. *wisteriana*, *C*. *maculata*), rather than the expected relationships recovered in previous work as depicted in Figure 6.

**Figure 6.**
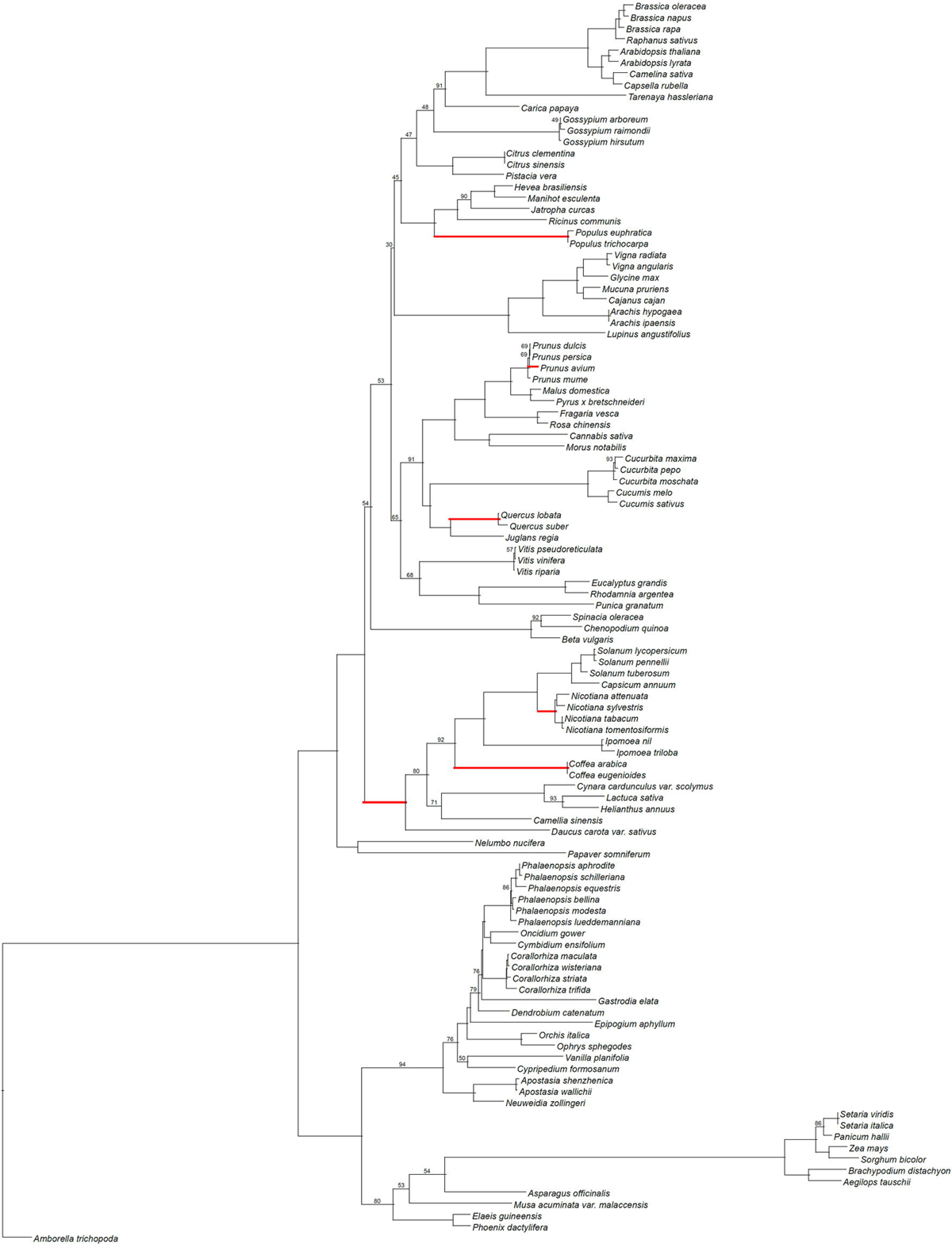
Maximum likelihood inferred topology of *WHY1* evolution with support values below 95 derived from 1,000 non-parametric bootstrap replicates, shown. Lineages highlighted in red represent those for which statistically-significant signal of episodic diversifying selection of *WHY1* was detected by the adaptive branch-site random effects likelihood test.

### Differential Expression

Canonical isoforms of *WHY1* were most highly expressed in all four species (see Table XX), but statistically significant elevated expression in aboveground tissues relative to those belowground was only detected in *C*. *trifida*. Gene-level expression of *WHY1* across both tissues was highest for *C*. *wisteriana* (244.5) and *trifida* (175.4) and lowest for *C*. *maculata* (131.9) and *striata* (95.3).

Expression of the canonical isoform of *WHY1* in *C. trifida* ranged from 4.15 to 9.64 TMM in belowground tissues (median = 7.17 TMM) and from 31.353 to 56.389 TMM in the aboveground tissues (median = 43.273 TMM). The log fold-change value was 2.56 between belowground and aboveground tissues for the canonical isoform of *C. trifida*, with a P value of 0.00018 and a false discovery rate (FDR) of 0.00374. Expression of the *WHY1* nearly canonical isoform in *C. striata* ranged from 5.547 to 16.859 TMM in belowground tissues (median = 12.31 TMM) and 17.28 and 30.538 TMM in aboveground tissues (median = 23.909 TMM). The *C. striata* canonical *WHY1* isoform had a log fold-change value of 0.396 between belowground and aboveground tissues, with a P value of 0.69 and FDR of 0.91. Expression of the canonical isoform of *WHY1* in *C. wisteriana* ranged from 9.256 to 23.023 TMM in belowground tissues (median = 15.663 TMM) and 34.792 to 74.955 TMM in aboveground tissues (median = 60.80 TMM). The *C. wisteriana* canonical isoform had a log fold-change value of 1.96 between belowground and aboveground tissues, with P value of 0.02 and FDR of 0.28. The expression of the canonical isoform in *C. maculata* had values of 11.064 and 13.947 TMM in belowground tissues (median = 12.51 TMM) and ranged from 26.179 to 45.999 TMM in aboveground tissues (median = 33.01 TMM). The *C. maculata* canonical isoform had a log fold-change value of 1.17 between belowground and aboveground tissues, with P value of 0.16 and FDR of 0.62.

The expression of non-canonical isoforms assembled from the read pools of all four species varied across tissues and samples, with some tissues or individuals not expressing splicing variants, and were not statistically different between the two tissue types in any of the four species. The *C*. *trifida* non-canonical isoform was expressed at relatively low levels across all aboveground and belowground tissues, the TMM of which ranged from 1.02 belowground to 4.374 aboveground.

Contrastingly, the TMM of the *C. wisteriana* non-canonical isoform ranged from 0 to 2.017 and was not detected in two of the belowground replicates for this species. The noncanonical *C*. *maculata* and *C. striata* isoforms were sporadically expressed across tissue types, with the TMM values of one isoform of *C*. *maculata* ranging from 0 to 3.564 and those of *C*. *striata* ranging from 0 to 4.471. Even in the case of the *C*. *striata* isoform for which the TMM value was 4.471 in one belowground sample, the value was either zero or less than 1 in the remainder of samples.

### Selection Analyses

The RELAX test inferred statistically significant signal for relaxation of selection pressure for the following test branch sets: *Corallorhiza* + *E*. *aphyllum* + *G*. *elata* (p = 0.001); *C*. *maculata* + *C*. *striata* + *E*. *aphyllum* + *G*. *elata* (p = 0.002); and *E*. *aphyllum* + *G*. *elata* (p = 0.001). However, runs of RELAX analyzing test branch sets comprising only *Corallorhiza* (p = 0.30) or *C*. *maculata* + *C*. *striata* (p = 0.62) did not infer significant signal for selection relaxation.

The FUBAR analysis inferred that 254 of 373 codons were under pervasive purifying selection (posterior probability threshold ≥ 0.9) and that no codons were under pervasive diversifying selection. Likewise, the BUSTED analyses did not infer statistically significant signal of gene-wide episodic diversifying selection for any test branch set, where p-values ranged from p = 0.15 for *E*. *aphyllum* + *G*. *elata* to p = 0.50 for *Corallorhiza*.

The aBSREL analysis recovered signal of episodic diversifying selection in seven of 217 branches in the *WHY1* tree (Figure 6) but did not infer statistically significant signal in any test branch set during analyses conducted in *a priori* mode. The terminal node leading to *Sorghum bicolor* (p = 2.76 x 10^-6^) was the only branch in the monocots that showed significant signal of episodic diversifying selection. Within the eudicots, the branches leading to *Prunus avium* (p = 2.5 x10^-5^), the *Populus* clade (p = 1.17 x 10^-7^), the *Quercus* clade (2.35 x 10^-3^), the *Nicotiana* clade (p = 5.42 x 10^-3^), the *Coffea* clade (6.96 x 10^-5^), and the asterid clade (p = 3.7 x 10-4) were inferred to contain signal of episodic diversifying selection.

## 4 Discussion

Our work represents the largest-scale investigation of *WHY1* evolution to date and reveals strong phylogenetic signal for this gene across multiple taxonomic levels in the angiosperms. Phylogenetic methods have emerged as the gold-standard for the inference of gene orthology (Münster et al., 1997; Emms and Kelly, 2015; Emms and Kelly, 2019) by which hundreds of genes have been determined to not only be highly conserved across plant lineages, but to exist as low- or single-copy (Wu et al., 2006; Duarte et al., 2010; De Smet et al., 2013). The inferred evolutionary relationships of *WHY1* (Figure 6) were largely congruent with our contemporary understanding of angiosperm phylogenetic relationships and recent inferences based on genomic-scale datasets (Angiosperm Phylogeny Group, 2016; Guo et al., 2021; Zhao et al., 2021). Likewise, the inferred relationships within the Orchidaceae were largely consistent with those inferred in other studies (e.g., Givnish et al., 2016; Pérez-Escobar et al., 2021; Serna-Sánchez et al., 2021), apart from *D*. *catenatum*, *G*. *elata*, and *E*. *aphyllum* forming an early-diverging, paraphyletic grade within the Epidendroids. However, our recovery of *C*. *trifida* as sister to the remainder of the species sampled from *Corallorhiza*, rather than *C*. *striata*, was not congruent with relationships inferred previously on the basis of other loci and even genomic-scale datasets (Barrett and Freudenstein, 2008; Barrett et al., 2014; Barrett et al., 2018). We are not surprised to infer discordance between the evolutionary history of *WHY1* and that of the phylogenetic history of *Corallorhiza*, since it is accepted that the evolutionary history of any gene can differ from that of the genome within which it is found (Pamilo and Nei, 1988; Maddison, 1997). That said, the recovery of the *Corallorhiza* species with the most intact plastome as sister to those with more degraded plastomes may indicate that the gene-species tree discordance we infer is due to functional convergence in *WHY1* sequence. For example, the phenylalanine substitution at position 336 shared by all *Corallorhiza* species aside from *C*. *trifida* may affect protein conformation and therefore *WHY1* function. Our findings of general congruence between the evolutionary history of *WHY1* across angiosperms supports the gene as single- or low-copy across the 110 angiosperm taxa sampled.

Differences in the degree of nucleotide conservation were evident amongst *WHY1* exons, with the transit peptide region consistently the most divergent throughout the lineages sampled. The diversity and evolution of transit peptides, and the apparent discrepancy between our perception of their functional importance and high sequence divergence, have long been of interest (Bruce, 2001; Patron and Waller, 2007; Christian et al., 2020). While the functional nature of a transit peptide might lead to an expectation of conservatism, low sequence similarity and patterns of mutation that we describe in *WHY1* are emerging as generalizable properties of plant transit peptides. For example, Christian et al. (2020) found that the transit peptides in the genomes of 15 genera sampled throughout the angiosperms had a mean pairwise percent identity of just 37.9% and that random indels drive transit peptide evolution. Our results provide evidence that the transit peptide of *WHY1* evolves similarly, with a pairwise percent identity of 32.5% across the angiosperms, and with evident substitutions and indels downstream of a homologous start codon being the most likely drivers of divergence in the gene region. In contrast, the portion of the chain encoding the ssDNA binding motif was the most conserved with a mean percent pairwise similarity of 72.1%. The only sizable indel observed in the gene region was a seven-codon insertion in *E*. *aphyllum*, a late-stage mycoheterotrophic orchid. It is likely that *E*. *aphyllum WHY1* results in a non-functional product, given that it encodes a premature stop codon that would result in a protein 70 amino acids shorter than that of any other sampled species. Interestingly, *WHY1* was more conserved overall at the nucleotide level amongst *Corallorhiza* species than were either expression patterns across tissues or exon inclusion in sequenced mRNAs across the trophic gradient. Amongst autotrophic species, nucleotide divergence of *WHY1* is particularly pronounced in the grasses, with the mean pairwise percent identity amongst all members of that clade versus the remainder of samples being more than 5% lower than that of the same comparison made for either of the late-stage mycoheterotrophic species sampled. The high substitution rate we inferred for *WHY1* in the grasses is interesting because the plastomes of many species in the clade are known to contain inversions and structural heteroplasmy within individual plants, of which the latter has only recently been described from their relatives the Cyperaceae (Doyle et al., 1992; Lee et al., 2020). The contrasting levels of nucleotide conservation in the transit and chain portions of *WHY1* across lineages suggest that lineage-specific functions of the transit peptide sequence could be a fruitful line of investigation, especially given that Christian et al. (2020) identified bias in amino acid usage between the plastid transit peptide sequences of monocot and eudicot lineages.

Alignment of *WHY1* revealed lineage-specific substitutions at sites inferred to be involved in structural conformation in both poorly and highly conserved gene regions. For example, the transit regions of both late-stage mycoheterotrophic species contained indels with little homology with the remainder of samples, and the sequence of *E*. *aphyllum* contained a premature stop codon in the typically highly conserved chain region followed downstream by four autapomorphic amino substitutions in an eight amino acid span. We identified substitutions that could underlie functional change in each *Corallorhiza* species, the most interesting of which was a substitution involving an alpha helix inferred in previous work conducted in *A*. *thaliana* (Cappadocia et al., 2013). The phylogenetic framework provided by our analysis suggests that the plesiomorphic codon state of position 336 was a leucine, which was substituted to an isoleucine in *C*. *trifida* and to a phenylalanine in the remainder of *Corallorhiza* sampled. We infer that the phenylalanine substitution in *Corallorhiza* species aside from *C*. *trifida* would result in the introduction of a benzene ring in which there were previously only aliphatic hydrocarbons. While our work revealed many nonsynonymous substitutions throughout *Corallorhiza* sequences, we did not identify nucleotide substitutions that obviously result in loss of function in these transitional mycoheterotrophic species.

Statistically significant shifts of selective regime were detected for late-stage mycoheterotrophic and some non-orchid autotrophic lineages. Our inference that 68.1% of *WHY1* codons are under pervasive purifying selection and none are under pervasive positive selection, despite the relatively low levels of nucleotide conservatism we documented, are congruent with the critical function of the gene across the angiosperms as has been documented in model and agricultural species in previous work. While we inferred disproportionately high substitution rates for *WHY1* in the grasses, we did not find evidence that divergence of those sequences was associated with diversifying selection. Despite the lack of pervasive positive selection in *WHY1* across the angiosperms, our identification of significant episodic diversifying selection in seven of 100 autotrophic lineages sampled suggests that a lineage-specific adaptive role of *WHY1* may be relatively common, though similar signal was not identified in sampled mycoheterotrophs. The lack of diversifying signal in any of our mycoheterotrophic species does not support neofunctionalization of the gene, which we find interesting given the myriad functions of *WHY1* and the diversifying signal identified in lineages across the angiosperms. Our findings together suggest that relaxation of *WHY1* selective constraint occurs after the transition to fully mycoheterotrophy, as significant signal of relaxed selection was only detected in branch sets containing *E*. *aphyllum* and *G*. *elata*, two late-stage fully mycoheterotrophic orchids.

Our analyses reveal that the expression of *WHY1* in *Corallorhiza* can differ by both tissue type and across the mycoheterotrophic gradient. Multiple studies have induced and characterized the effects of differential expression of *WHY1* by exposure to biotic and abiotic stimuli, implicating roles for the gene ranging from mediating drought stress (Ruan et al., 2022) to pathogen response (Desveaux et al., 2000; Sun et al., 2023). A minimum level of *WHY1* expression could be expected due to roles of *WHY1* that are not involved in photosynthesis, such as the maintenance of telomere length of nuclear chromosomes (Yoo et al., 2007). Our analyses are the first to characterize *WHY1* expression in non-model or non-cultivated plant species, and therefore baseline expectations for tissue-specific expression levels for wild species have not yet been established. However, the values observed in *Corallorhiza* are within the expressive ranges reported in studies of *A*. *thaliana* (4-76 TPM; Liu, et al., 2012; Mergner et al., 2020) and *Solanum tuberosum* (7-50 TPM; The Potato Genome Sequencing Consortium, 2011). Additionally, the Klepikova *Arabidopsis* Atlas (Klepikova et al., 2016) and 1,122 tissue-specific samples available via the *Arabidopsis* RNA-seq Database (http://ipf.sustech.edu.cn/pub/athrdb/; accessed 8 March 2023) evidence that *WHY1* expression should be expected to be lower in roots or rhizomes than in leaves, which is congruent with expression patterns in *C*. *trifida*. Previous studies reporting *WHY1* expression have been conducted in species with larger individuals with typical, non-reduced morphologies allowing for finer-scale investigations of tissue-specific expression than can be conducted in *Corallorhiza*, due to a lack of leaf laminae and roots in the latter. However, the patterns of expression we found across *Corallorhiza* tissues are congruent with those known for *WHY1*. While gene-level expression across tissues was highest for *C*. *wisteriana* and *trifida*, tissue-specific expression was only statistically significant in *C*. *trifida*, after correcting for repeated testing. Our findings characterizing similar *WHY1* expression amongst the belowground tissues of *Corallorhiza* provide for a hypothesized minimum expression level of canonical *WHY1*, which is similar to aboveground levels of expression in *Corallorhiza* species aside from *C*. *trifida.* Taken together, our results provide evidence for a trajectory beginning with differential expression of *WHY1* between aboveground and belowground tissues of the most photosynthetically capable *Corallorhiza* species to similar expression levels between above and belowground tissues of the latest-stage mycoheterotrophic members of the genus. These results suggest that downregulation of *WHY1* in aboveground tissues is correlated with increased mycoheterotrophy, but that it is unlikely to underlie the transition to mycoheterotrophy since the tissue-level expression patterns of the gene in *C*. *trifida* remain like those described from autotrophic plants.

Our work is the first to characterize alternative splicing of, and intron retention in, *WHY1.* Approximately 70% of plant genes with multiple exons can be expected to be alternatively spliced (Chamala et al., 2015), and intron retention is a common form of alternative splicing in plants (Ner-Gaon et al., 2004). Our finding of intron retention in one isoform from each *Corallorhiza* species aside from *C*. *trifida* is the first such event described for the gene. It has long been recognized that intron retention is most common in transcripts of genes like *WHY1* which serve roles related to photosynthesis and stress response (Ner-Gaon et al., 2004), a finding also supported by work investigating the effects of plant stressors on levels of alternative splicing (Filichkin et al., 2018; Jabre et al., 2019). In fact, tissue-specific differential intron retention has been shown to be an inducible stress response in *Populus trichocarpa* (Filichkin et al., 2018). Modification of *WHY1*, including inserted sequence, has long been used to study the effects of mutations on the function of the gene and to induce knockouts (Desveaux et al., 2004; Yoo et al., 2007; Maréchal et al., 2009), work which helped identify the many pathways that *WHY1* is involved in. For example, Yoo et al., (2007) and Maréchal et al. (2009) leveraged knockouts caused by T-DNA insertions into *WHY1* to reveal the critical role the gene plays in maintaining telomere length and plastome stability in *Arabidopsis thaliana*, respectively. Similarly, Prikryl et al. (2008) and Qui et al., (2022) found that double knockout *WHY1* mutants were characterized by a lethal albino phenotype after the development of a few leaves, with Qui et al. (2022) also characterizing divergent splicing and mRNA editing of plastid genes in *WHY1* mutants. Our discovery of intron-retaining transcripts from all *Corallorhiza* species aside from *C*. *trifida*, and more transcriptional isoforms in *Corallorhiza* species in later stages of mycoheterotrophy, reveals a negative correlation between increased splicing variation and both plastome stability and chlorophyll concentration in tissues (Barrett et al., 2014).

We find it unlikely that the intron-containing isoforms result in functional products, since a premature stop codon results from intron retention. The recovery of non-canonical and intron-retaining *WHY1* isoforms across individuals and tissues of *Corallorhiza* could be signal of idiosyncratic spliceosome regulation in mycoheterotrophic species, epitranscriptomic differences (Jabre et al., 2019), differential responses to stress in sampled tissues (Filichkin et al., 2018), or the expression of multiple, divergent copies of the gene. However, we propose that reduced fidelity in the spliceosome of a mycoheterotrophic plant is the most likely cause of the observed splicing variation, given the phylogenetic and sequencing data at hand. We predict that changes in the expression and splicing of *WHY1* across *Corallorhiza* are likely due to the alteration of one or more pathways involved in gene regulation, since we did find nucleotide-level changes that could be responsible.

Our work is the first to characterize the evolution of a transcription factor that could impact the genetic and phenotypic changes that occur along the path to full mycoheterotrophy. The roles that *WHY1* plays in plastome stability (Parent et al., 2011; Lepage, 2013), defense responses (Lin et al., 2020), and leaf senescence (Lin et al., 2019), together position the gene as a worthwhile target for the study of the molecular underpinnings of the transition to mycoheterotrophy. Heterotrophy in plants is associated with genomic restructuring, where a trend of plastome contraction and nuclear genome expansion via rampant repetitive element accumulation has commonly been observed (Barrett et al., 2014; Lyko and Wicke, 2021). Dramatic reductions of plastome length and gene content of mycoheterotrophic plants have been documented, ranging from minimal degradation in early-transitional orchids such as *Corallorhiza* (Barrett and Davis, 2012) to pronounced degradation in late-transitional orchids such as *Epipogium* and *Pogoniopsis* (Schelkunov et al., 2015; Klimpert et al., 2022). Our finding of putatively functional but increasingly alternatively spliced *WHY1* across the *Corallorhiza* trophic gradient is not necessarily surprising, given the minimally destabilized plastomes of the group. For example, Barrett and Davis (2012) found that the plastome of *C*. *striata*, the most destabilized of the *Corallorhiza* species included here, is only about 6% reduced relative to that of a leafy, autotrophic relative. Despite their relatively intact states, *Corallorhiza* plastomes are in various stages of degradation (Barrett et al., 2014), and our work here reveals a negative correlation between both increased putatively aberrant splicing and nucleotide-level divergence of *WHY1* with plastome stability across the sampled species. Likewise, the plastomes of *Gastrodia elata* and *Epipogium aphyllum*, both late-stage mycoheterotrophs for which our analyses show that *WHY1* contains premature stop codons and significant signal of relaxed selection, are both extremely reduced and syntenically disrupted (Yuan et al., 2018; Chen et al., 2020; Xu et al., 2021). Taken together, our findings provide evidence of a negative correlation between increased divergence in sequence, splicing of *WHY1*, and plastome stability in early- to late-stage mycoheterotrophic orchids.

## 5 Conclusions

Our work showcases the rich opportunities afforded by mycoheterotrophic plants not just for the study of the evolution of *WHY1* but for any gene of which homozygous knockouts can result in a fatal phenotype in autotrophic plants. Continued investigation of non-autotrophic plant lineages promises to fill gaps in our understanding of the precursors and consequences of genomic instability, and even the minimum gene space of land plants. Here we presented findings of non-synonymous nucleotide substitutions in functionally annotated regions in *Corallorhiza WHY1* sequence, and a high degree of divergence in *WHY1* in late-stage fully mycoheterotrophic orchids. However, our results together suggest that changes to the expression and splicing of *WHY1* occur prior to the establishment of obviously deleterious genomic substitutions that would render the TF non-functional in late stage mycoheterotrophic orchids. In sum, our work supports a correlation between decreased expression and increased alternative splicing of *WHY1* with plastome degradation in a group of early-transitional mycoheterotrophic orchids but does not implicate divergent *WHY1* function as a primary factor in the transition from partial to full mycoheterotrophy.

## 7 Author Contributions

BS and CB contributed to the conception and design of the study. CB collected *Corallorhiza* samples for RNAseq. RM and BS collected all publicly available samples, conducted wet-lab work, Nanopore sequencing, and bioinformatic analyses. RM wrote the first manuscript draft. All authors contributed to manuscript revision and have read and approved of the final version.

## Acknowledgments

We would like to acknowledge John V. Freudenstein for supplying DNAs used for Nanopore sequencing, the West Virginia University Genomics Core Facility for their services and technical expertise. We also thank the USDA Forest Service and California State Parks for permission to collect material.

## 9 Data Availability Statement

Reads resulting from RNAseq and Nanopore sequencing are available via NCBI (BioProject PRJNA984634; released upon acceptance) as components of BAM formatted files where reads are mapped to the assembled transcript or contig. All sequence alignments analyzed are available in FASTA format via a public GitHub repository (https://github.com/btsinn/whirly1evolution.git; archived via Zenodo upon acceptance).

---

**Table 1.** *In-silico* expression of *WHY1* across species, tissues, and biological replicates. The first row for a species contains values for gene-level expression, while subsequent rows contain values for a specific isoform. Trinity isoform and biological replicate identifiers are provided in Table S3. Expression values are trimmed-mean-of-means-transformed (TMM) transcripts per million (TPM) in order to maintain comparability amongst biological replicates. An * denotes a canonical isoform while ^ψ^ denotes an isoform with a retained intron. Values shown have been truncated, TMM to the tenth, LogFC to the hundredth, and P- and Q-values to the thousandth.

| Species<br>Gene or Isoform | Belowground Tissue<br>(Biological Replicate #) |  |  | Aboveground Tissue<br>(Biological Replicate #) |  |  | LogFC | P-value<br>(Q-value) |
| --- | --- | --- | --- | --- | --- | --- | --- | --- |
| *C. trifida* | 1 | 2 | 3 | 1 | 2 | 3 |  |  |
| Gene | 12.7 | 11.6 | 4.7 | 31.4 | 71.0 | 43.7 | 2.12 | 0.004<br>(0.057) |
| Isoform 1 | 5.0 | 1.8 | 1.0 | 1.4 | 2.9 | 4.3 | 0.12 | 0.915<br>(0.998) |
| Isoform 2* | 7.7 | 9.6 | 4.1 | 31.3 | 56.3 | 42.0 | 2.56 | 0.0001<br>(0.003) |
| *C. wisteriana* | 1 | 2 | 3 | 1 | 2 | 3 |  |  |
| Gene | 14.2 | 23.0 | 8.3 | 84.3 | 76.7 | 37.8 | 1.91 | 0.019<br>(0.176) |
| Isoform 1<sup>ψ</sup> | 0 | 0.8 | 0 | 1.0 | 1 | 2.0 | 2.36 | 0.313<br>(1) |
| Isoform 2* | 14.7 | 23.0 | 9.2 | 74.9 | 72.6 | 34.7 | 1.96 | 0.021<br>(0.283) |
| *C. maculata* | 1 | 2 | NA | 1 | 2 | 3 |  |  |
| Gene | 16.6 | 12.7 |  | 47.0 | 25.3 | 30.0 | 1.17 | 0.162<br>(0.624) |
| Isoform 1<sup>ψ</sup> | 2.0 | 1.8 |  | 0.5 | 0 | 3.5 | -0.51 | 0.746<br>(0.970) |
| Isoform 2* | 13.9 | 11.0 |  | 45.9 | 26.1 | 26.8 | 1.33 | 0.119<br>(0.387) |
| *C. striata* | 1 | 2 | 3 | 1 | 2 | NA |  |  |
| Gene | 21.3 | 16.4 | 7.8 | 32.1 | 17.3 |  | 0.62 | 0.525<br>(0.804) |
| Isoform 1 | 0.9 | 0 | 2.3 | 0 | 0 |  | NA | NA |
| Isoform 2 | 16.8 | 14.5 | 5.5 | 30.5 | 17.2 |  | 0.39 | 0.695<br>(0.918) |
| Isoform 3<sup>ψ</sup> | 4.4 | 0.7 | 0.5 | 0.8 | 0 |  | -2.88 | 0.254<br>(0.536) |

