## supplementary materials for "A case study of *Whirly1* (*WHY1*) evolution in the angiosperms: altered sequence, splicing, and expression in a clade of early-transitional mycoheterotrophic orchids"

### **Supplemental Information**

**Table S1**

*Angiosperm WHY1 alignment accession*

| **Order** | **Family** | **Taxon** | **Accession** |
| --- | --- | --- | --- |
| Poales | Poaceae | *Aegilops tauschii subsp. tauschii* | XM_020331194 |
| Amborellales | Amborellaceae | *Amborella trichopoda* | XM_006842663 |
| Asparagales | Orchidaceae | *Apostasia shenzhenica* | PKA64494.1 |
| Asparagales | Orchidaceae | *Apostasia wallichii* | AUTC011479 |
| Asparagales | Orchidaceae | *Apostasia zollingeri* | NZTC028581 |
| Brassicales | Brassicaceae | *Arabidopsis lyrata subsp. lyrata* | XM_021013506 |
| Brassicales | Brassicaceae | *Arabidopsis thaliana* | NM_101308 |
| Fabales | Fabaceae | *Arachis hypogaea* | XM_025783221 |
| Fabales | Fabaceae | *Arachis ipaensis* | XM_016335481 |
| Asparagales | Asparagaceae | *Asparagus officinalis* | XM_020395017 |
| Caryophyllales | Amaranthaceae | *Beta vulgaris subsp. vulgaris* | XM_010684944 |
| Poales | Poaceae | *Brachypodium distachyon* | XM_003557150 |
| Brassicales | Brassicaceae | *Brassica napus* | XM_013822335 |
| Brassicales | Brassicaceae | *Brassica oleracea var. oleracea* | XM_013746574 |
| Brassicales | Brassicaceae | *Brassica rapa* | XM_009112295 |
| Fabales | Fabaceae | *Cajanus cajan* | XM_020380279 |
| Brassicales | Brassicaeae | *Camelina sativa* | XM_010497756 |
| Ericales | Theaceae | *Camellia sinensis* | XM_028229881 |
| Rosales | Cannabaceae | *Cannabis sativa* | XM_030628788 |
| Brassicales | Brassicaeae | *Capsella rubella* | XM_006303212 |
| Solanales | Solanaceae | *Capsicum annuum* | XM_016693060 |
| Brassicales | Caricaeae | *Carica papaya* | XM_022041728 |
| Caryophyllales | Amaranthaceae | *Chenopodium quinoa* | XM_021890501 |
| Sapindales | Rutaceae | *Citrus clementina* | XM_006438564 |
| Sapindales | Rutaceae | *Citrus sinensis* | XM_006483162 |
| Gentianales | Rubiaceae | *Coffea arabica* | XM_027242545 |
| Gentianales | Rubiaceae | *Coffea eugenioides* | XM_027295752 |
| Asparagales | Orchidaceae | *Corallorhiza maculata* | TRINITY DN55535 c1 g1 i2 |
| Asparagales | Orchidaceae | *Corallorhiza striata* | TRINITY DN2919 c0 g1 i2 |
| Asparagales | Orchidaceae | *Corallorhiza trifida* | TRINITY DN3392 c0 g1 i2 |
| Asparagales | Orchidaceae | *Corallorhiza wisteriana* | TRINITY DN2856 c1 g1 i2 |
| Cucurbitales | Cucurbitaceae | *Cucumis melo* | XM_008458246 |
| Cucurbitales | Cucurbitaceae | *Cucumis sativus* | XM_011659181.2 |
| Cucurbitales | Cucurbitaceae | *Cucurbita maxima* | XM_023143979 |
| Cucurbitales | Cucurbitaceae | *Cucurbita moschata* | XM_023091003 |
| Cucurbitales | Cucurbitaceae | *Cucurbita pepo subsp. pepo* | XM_023690277 |
| Asparagales | Orchidaceae | *Cymbidium ensifolium* | CETC024278 |
| Asterales | Asteraceae | *Cynara cardunculus var. scolymus* | XM_025138614 |
| Asparagales | Orchidaceae | *Cypripedium formosanum* | CFTC020472 |
| Apiales | Apiaceae | *Daucus carota subsp. sativus* | XM_017376908 |
| Asparagales | Orchidaceae | *Dendrobium catenatum* | XM_020824140 |
| Arecales | Arecaceae | *Elaeis guineensis* | XM_010916845 |
| Asparagales | Orchidaceae | *Epipogium aphyllum* | GFWK01035004.1 |
| Myrtales | Myrtaceae | *Eucalyptus grandis* | XM_010068645 |
| Rosales | Rosaceae | *Fragaria vesca subsp. vesca* | XM_004297283 |
| Asparagales | Orchidaceae | *Gastrodia elata* | GETC018859 |
| Fabales | Fabaceae | *Glycine max* | XM_014779345 |
| Malvales | Malvaceae | *Gossypium arboreum* | XM_017764060 |
| Malvales | Malvaceae | *Gossypium hirsutum* | XM_016819285 |
| Malvales | Malvaceae | *Gossypium raimondii* | XM_012607821 |
| Asterales | Asteraceae | *Helianthus annuus* | XM_022156146 |
| Malpighiales | Euphorbiaceae | *Hevea brasiliensis* | XM_021809249 |
| Solanales | Convolvulaceae | *Ipomoea nil* | XM_019309367 |
| Solanales | Convolvulaceae | *Ipomoea triloba* | XM_031244589 |
| Malpighiales | Euphorbiaceae | *Jatropha curcas* | XM_012236542 |
| Fagales | Juglandaceae | *Juglans regia* | XM_018957158 |
| Asterales | Asteraceae | *Lactuca sativa* | XM_023887217 |
| Fabales | Fabaceae | *Lupinus angustifolius* | XM_019599127 |
| Rosales | Rosaceae | *Malus domestica* | NM_001294042 |
| Malpighiales | Euphorbiaceae | *Manihot esculenta* | XM_021747739 |
| Cucurbitales | Cucurbitaceae | *Momordica charantia* | XM_022295640 |
| Rosales | Moraceae | *Morus notabilis* | XM_024165601 |
| Fabales | Fabaceae | *Mucuna pruriens* | RDX92873.1 |
| Zingiberales | Musaceae | *Musa acuminata subsp. malaccensis* | XM_009382162 |
| Solanales | Solanaceae | *Nicotiana attenuata* | XM_019401472 |
| Solanales | Solanaceae | *Nicotiana sylvestris* | XM_009760051 |
| Solanales | Solanaceae | *Nicotiana tabacum* | XM_016618964 |
| Solanales | Solanaceae | *Nicotiana tomentosiformis* | XM_009600939 |
| Asparagales | Orchidaceae | *Oncidium gower* | OGTC013046 |
| Asparagales | Orchidaceae | *Ophrys sphegodes* | OSTC004794 |
| Asparagales | Orchidaceae | *Orchis italica* | OITC019736 |
| Poales | Poaceae | *Oryza sativa* | XM_015785850 |
| Poales | Poaceae | *Panicum hallii* | XM_025954601 |
| Asparagales | Orchidaceae | *Phalaenopsis aphrodite* | PATC126989 |
| Asparagales | Orchidaceae | *Phalaenopsis bellina* | PBTC020526 |
| Asparagales | Orchidaceae | *Phalaenopsis equestris* | PETC023606 |
| Asparagales | Orchidaceae | *Phalaenopsis lueddemanniana* | PLTC001961 |
| Asparagales | Orchidaceae | *Phalaenopsis modesta* | PMTC004074 |
| Asparagales | Orchidaceae | *Phalaenopsis schilleriana* | PSTC007514 |
| Arecales | Arecaceae | *Phoenix dactylifera* | XM_026810121 |
| Sapindales | Anacardiaceae | *Pistacia vera* | XM_031393836 |
| Malpighiales | Salicaceae | *Populus euphratica* | XM_011027665 |
| Malpighiales | Salicaceae | *Populus trichocarpa* | XM_002311565 |
| Rosales | Rosaceae | *Prunus avium* | XM_021951697 |
| Rosales | Rosaceae | *Prunus dulcis* | XM_034343292 |
| Rosales | Rosaceae | *Prunus mume* | XM_008223440 |
| Rosales | Rosaceae | *Prunus persica* | XM_007222413 |
| Myrtales | Lythraceae | *Punica granatum* | XM_031546949 |
| Rosales | Rosaceae | *Pyrus x bretschneideri* | XM_009362111 |
| Fagales | Fagaceae | *Quercus lobata* | XM_031100709 |
| Fagales | Fagaceae | *Quercus suber* | XM_024072291 |
| Brassicales | Brassicaceae | *Raphanus sativus* | XM_018592404 |
| Myrtales | Myrtaceae | *Rhodamnia argentea* | XM_030687962 |
| Malpighiales | Euphorbiaceae | *Ricinus communis* | XM_002520082 |
| Rosales | Rosaceae | *Rosa chinensis* | XM_024338948 |
| Poales | Poaceae | *Setaria italica* | XM_004964480 |
| Poales | Poaceae | *Setaria viridis* | XM_034736473 |
| Solanales | Solanaceae | *Solanum lycopersicum* | NM_001302900 |
| Solanales | Solanaceae | *Solanum pennellii* | XM_015219096 |
| Solanales | Solanaceae | *Solanum tuberosum* | NM_001288226 |
| Poales | Poaceae | *Sorghum bicolor* | XM_002436422 |
| Caryophyllales | Amaranthaceae | *Spinacia oleracea* | XM_021988930 |
| Brassicales | Capparaceae | *Tarenaya hassleriana* | XM_010553210 |
| Asparagales | Orchidaceae | *Vanilla planifolia* | VPTC010179 |
| Fabales | Fabaceae | *Vigna angularis* | XM_017569082 |
| Fabales | Fabaceae | *Vigna radiata var. radiata* | XM_014651952 |
| Vitales | Vitaceae | *Vitis pseudoreticulata* | MN395402 |
| Vitales | Vitaceae | *Vitis riparia* | XM_034838923 |
| Vitales | Vitaceae | *Vitis vinifera* | XM_002277242 |
| Poales | Poaceae | *Zea mays* | NM_001130117 |

**Table S2.**

Oxford Nanopore consensus read lengths and mean coverages.

| Species (collection number) | Number of Oxford Nanopore Reads | consensus sequence length (bp) | mean coverage |
| --- | --- | --- | --- |
| *Corallorhiza trifida* (2227) | 25,546 | 396 | 4,034.9 |
| *Corallorhiza maculata* (2123) | 74,261 | 382 | 8,002.5 |
| *Corallorhiza striata* (2084) | 61,810 | 381 | 10,617.6 |

**Table S3.**

Trinity transcript identifiers and biological replicate source information corresponding to Table 1.

| **Species** | **Trinity Transcript Identifer** | **Belowground Tissue**  **Sample Identifiers** | **Aboveground Tissue**  **Sample Identifiers** |
| --- | --- | --- | --- |
| **C. trifida** |  | | |
| Isoform 1 | TRINITY_DN3392_c0_g1_i1 | **16_4R_A**  **16_4R_B**  **CFB** | **16_4F_A**  **16_4F_B**  **CFB** |
| Isoform 2 | TRINITY_DN3392_c0_g1_i2 |  |  |
| **C. wisteriana** |  | | |
| Isoform 1 | TRINITY_DN2856_c1_g1_i1 | **16_3R_A**  **16_3R_B**  **CFB** | **16_3F_A**  **16_3F_B**  **CFB** |
| Isoform 2 | TRINITY_DN2856_c1_g1_i2 |  |  |
| **C. maculata** |  | | |
| Isoform 1 | TRINITY_DN55535_c1_g1_i1 | **16_1macR**  **CFB** | **16_1F**  **437**  **CFB** |
| Isoform 2 | TRINITY_DN55535_c1_g1_i2 |  |  |
| **C. striata** |  | | |
| Isoform 1 | TRINITY_DN2919_c0_g1_i1 | **16_2R_A**  **16_2R_B**  **CFB** | **STR_2F**  **CFB** |
| Isoform 2 | TRINITY_DN2919_c0_g1_i2 |  |  |
| Isoform 3 | TRINITY_DN2919_c0_g1_i4 |  |  |
